# Empowering human-like walking with a bio-inspired gait controller for an under-actuated torque-driven human model

**DOI:** 10.1101/2023.12.10.571012

**Authors:** Samane Amini, Iman Kardan, Ajay Seth, Alireza Akbarzadeh

## Abstract

Human gait simulation plays a crucial role in providing insights into various aspects of locomotion, such as diagnosing injuries and impairments, assessing abnormal gait patterns, and developing assistive and rehabilitation technologies. To achieve more realistic results in gait simulation, it is necessary to utilize a comprehensive model that closely replicates the kinematics and kinetics of the human gait pattern. OpenSim software provides anthropomorphic and anatomically accurate human skeletal structures that enable users to create personalized models for individuals to accurately replicate real human behavior. However, torque-driven models face challenges in balancing unactuated degrees of freedom during forward dynamic simulations. Adopting a bio-inspired strategy that ensures an individual’s balance with a minimized energy expenditure, this paper proposes a gait controller for a torque-deriven OpenSim model to achieve a stable walking. The proposed controller takes a model-based approach to calculate a “*Balance Equivalent Control Torque*” and uses the concept of the hip-ankle strategy to distribute this balance torque to the lower-limb joints. To optimize the controller gains and the “*Balance Distribution Coefficients*”, an interface is stablished between MATLAB and OpenSim that is capable of conducting controllable forward dynamic simulations. The simulation results demonstrate that the torque-driven model can walk naturally with joint torques suitably matching experimental data. The robustness of the bio-inspired gait controller is also assessed by applying a range of external forces on the upper body to disturb the model. The robustness analysis demonstrates the quick and effective balance recovery mechanism of the proposed bio-inspired controller.

## Introduction

Forward dynamic simulation is a computational technique widely used in biomechanical systems to accurately simulate human movements[1-3]. This approach enables the model to closely replicate human behavior, including dynamic variables such as forces, torques, and motion trajectories[4]. To simulate human movements, various models have been used so far, ranging from simple mechanical models with biped models with low degrees of freedom to neuromuscular models, each with different capabilities[5-8].The OpenSim musculoskeletal human model is widely utilized for accurately simulating human movement and studying biomechanics[9]. It offers a detailed representation of the human body, including the skeletal structure and muscles[10]. Moreover, OpenSim skeletal models hold greater significance compared to mechanical models due to their anthropomorphic and anatomically accurate structure, as well as their scalability using motion capture[11]. However, the most critical challenge of torque-driven models are the existence of unactuated degrees of freedom and the requirement to implement an algorithm to maintain balance[12, 13]. In fact, in the forward dynamic simulation of the skeletal model of a human in OpenSim, an unactuated joint, pelvis joint, is not directly controlled or actuated by muscles or external forces. Consequently, it is crucial to devise a method for maintaining the balance and stability of the model during dynamic simulations. Hence, our primary objective in this study is to integrate a reliable strategy that ensures human balance in the simulation. A considerable amount of research has been conducted on this area up to the present. One prevalent method for assessing and controlling balance during gait is the Zero-Moment-Point (ZMP) method[14-17]. This approach ensures the static and dynamic stability of the human model by controlling the ZMP to remain within the foot-support region, which is formed by connecting the foot-ground contact points. Another method involves determining appropriate initial conditions for the joints. In[18], the initial conditions for the stable limit cycle of the passive dynamic model are calculated based on the “basin of attraction” concept. By setting the initial conditions properly, stable walking cycles can be achieved, closely resembling natural human gait. However, when there are changes in the walking pattern or joint trajectory, it becomes necessary to update the initial conditions for the new motion.

Open-loop models that solely rely on appropriate initial conditions are not robust against external disturbances and environmental changes. To address this, several closed-loop techniques have been proposed to provide robust balance algorithms for biped models. These include momentum conservation [19, 20], phase resetting[18, 21], impedance control of joints[22-24], and intermittent control[25-28]. Nonetheless, designing and implementing a balance controller that accurately replicates human gait patterns with reliable torque/power in the joints remains an open and challenging area of research.

One of the key factors contributing to the analysis of human gait is the remarkable efficiency of the human body in minimizing energy expenditure while walking[29]. In fact, the human body adjusts its walking speed and stride length to optimize energy expenditure. Studies have consistently shown the existence of an optimal walking speed at which the metabolic cost is minimized[30-32]. Individuals tend to naturally select a walking speed close to this optimal value, which indicates the body’s innate ability to self-regulate and minimize energy expenditure.

In reality, achieving an accurate representation of human gait involves simulating the model in such a manner that either the metabolic cost (in the case of a musculoskeletal model) or the mechanical energy (in the context of a skeletal model) is minimized. As a result, our goal is to employ a suitable equilibrium technique that preserves human stabilization while concurrently minimizing mechanical energy expenditure. To attain this objective, we adopted the hip-ankle human balance strategy, which involves a reduced count of joints responsible for upholding model stability, consequently leading to a reduction in energy costs. According to the ankle-hip strategy, if a human is standing in an upright posture and a perturbation is applied, depending on the size of the disturbance, the person will engage one or both of the ankle and the hip joints or will take a step to prevent unbalancing[33, 34]. Furthermore, we developed a controllable platform by integrating OpenSim with the MATLAB environment to simulate the movement of an under-actuated human model. This interface allowed us to construct a forward dynamic simulation of the human model of OpenSim through MATLAB command script.

This paper is organized as follows: First, the gait controller of the under-actuated model is described. Second, the bio-inspired balance approach of the gait controller is explained. In the next section, the optimization process of the controller is described. Finally, the performance of the proposed controller is validated in the results and discussion section.

## Methodology

The human body has been modeled as a multibody system, formed by rigid bodies linked via ideal joints and controlled by torque actuators. This section elucidates the process of developing a human gait controller within the context of the multibody dynamics of the model. To achieve this, the equation of motion for the human model will be dissected into actuated and un-actuated components.

### Gait control of under-actuated model

Multibody dynamic equation of musculoskeletal human model is defined as follows[35]:

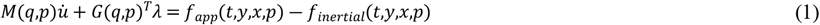

where *t,p, x*(*t*),*y*(*t*) stand for time, time-invariant parameters, control, and time-dependent states. *q*(*t*) and *u*(*t*) are generalized coordinate and generalized speed. In addition, *M,f*_*app*_, *f*_*inertial*_, and *λ*(*t*) are the mass matrix, the applied force involving gravity and muscle dynamics, the centripetal and Coriolis terms, and Lagrange multipliers respectively. To estimate the muscle forces underlying a given motion, some optimization methods are commonly used. These methods aim to find the muscle force distribution that satisfies certain criteria, such as minimizing the metabolic cost, matching experimental data, or maximizing task performance. However, estimating muscle forces in musculoskeletal models is a computationally intensive task due to the complexity of the underlying optimization problem.

Human torque-driven models are a computational representation of the human dynamic system that focuses on simulating and analyzing human movement primarily driven by joint torques. However, the most critical challenges of torque-driven models is the existence of unactuated degrees of freedom and the need for implementing an algorithm to maintain the balance. For a two-dimensional human model in sagittal plane, pelvis motions (one rotation and two vertical and horizontal translations) are the unactuated degrees-of-freedom that result in the model being underactuated and inherently unstable. The underactuated locomotion system consists of six degrees-of-freedom in each leg, including the hip, knee, and ankle joints. Additionally, it encompasses vertical, horizontal, and rotational movements in the upper body limb. The mathematical model derived by Lagrange law is simplified as,

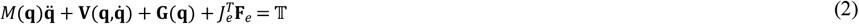

where, 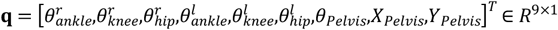 is the vector of joints variables and 𝕋 ∈ *R*^*9*×1^ is the vector of corresponding joint torques with zero torques/forces for pelvis DoFs. *M*(**q**) ∈ *R*^*9*×*9*^ is the mass matrix, 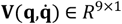 is the vector of centrifugal and Coriolis forces, **G** (**q**) ∈ *R*^*9*×1^ is the vector of gravitational forces, **F**_*e*_ ∈ *R*^*n*×1^ is the vector of external force components, ***J***_*e*_ ∈ *R*^*n*×*9*^ is a Jacobian matrix relating the position of the applied external forces to the joint variables, and *n* is the number of external force components, including ground reaction forces. Decomposing actuated and unactuated degrees-of-freedom, Eq. (2) may be re-written as,

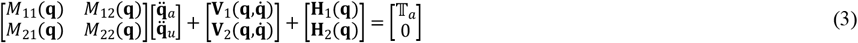

where, 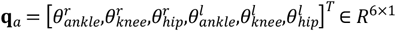 is the vector of actuated joints variables, 𝕋_*a*_ ∈ *R*^6×1^ is the vector of corresponding joint torques, **q**_*u*_ = [θ_*Pelvis*_,*X*_*Pelvis*_,*Y*_*Pelvis*_]^*T*^ ∈ *R*^3×1^ is the vector of unactuated degrees of freedom, and **H**_*i*_(**q**) ∈ *R*^*n*×1^ includes the gravitational and external forces/torques. The actuated and un-actuated dynamics are derived by dividing the overall system into two subsystems as,

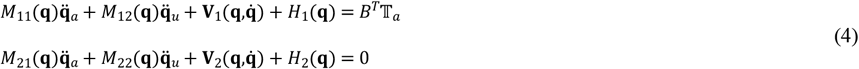

Thus, differential equations of two subsystems are obtained as,

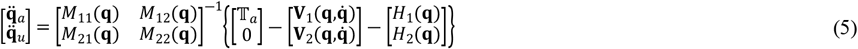

By simplifying the above equation, we have

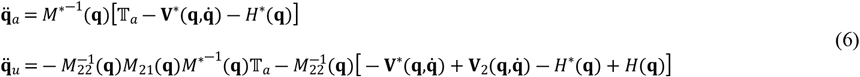

in which,

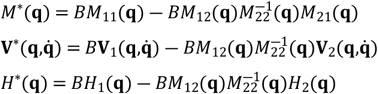

Therefore, the dynamic structure of the underactuated walking model is explained by the differential equation of two actuated and unactuated dynamics (6). Here, we define two equivalent controls for the actuated and un-actuated dynamics as 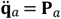 and 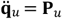 [36]. In this way, the joint moment control becomes:

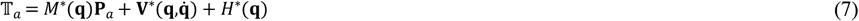

The primary objective is for the walking model to accurately follow a specified path trajectory while maintaining stability in the upper body. Hence, to track the path trajectory 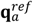 by the actuated joints **q**_*a*_,the equivalent control **P**_*a*_ is designed as,

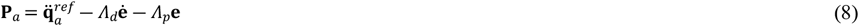

where 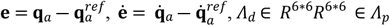, are positive diagonal matrices. Furthermore, to ensure stability of the unactuated subsystem, the equivalent control **P**_*u*_ is adjusted in accordance with the following equation:

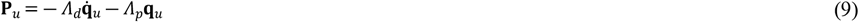

Ultimately, to achieve stabilization of the overall system, the final equivalent control for the underactuated system is generated by means of a linear combination, expressed as,

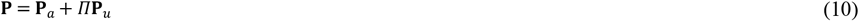

where *Π* represents a balancing distribution allocated to the actuated joints. By substituting **P** from equation (7) instead of **P**_*a*_ in equation (8), the joint moment control for balanced locomotion is computed as described in equation (11).

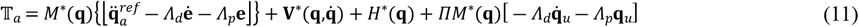

Through simplifying the moment equation, we have:

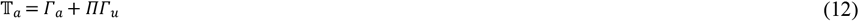

where,

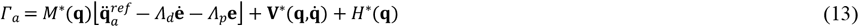

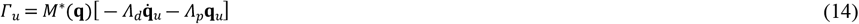

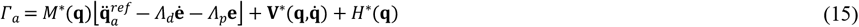

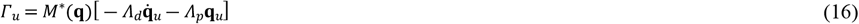

### Bio-inspired balance approach

According to Equation (12), the skeletal joint moment comprises both trajectory tracking control and balance control. The first component involves generating joint moments to enable the human model to follow a reference path, while the second component ensures the provision of torques necessary for maintaining balance during walking.

Based on our knowledge, the main characteristic of a normal gait, often referred to as an efficient or economical gait, is indeed the minimization of energy expenditure. In fact, when walking or running, the human body has evolved to optimize energy usage by employing various biomechanical strategies. Hence, the objective of this paper is to employ a biomechanical approach to design balance distribution coefficients that minimize the required locomotion energy. To achieve this goal, a method is adopted in which a smaller number of joints are involved in maintaining human balance, resulting in the production of less net balance momentum. The hip-ankle strategy, which is a neuromuscular control mechanism employed by humans to maintain balance during standing and walking, is used to preserve balance in mechanical biped models[37]. According to this strategy, when relatively small disturbances affect the center of mass (CoM), humans tend to employ the ankle strategy. In such cases, ankle torque is utilized to restore the CoM to its desired position and maintain balance. However, when faced with a significantly large disturbance, a torque is applied to the hip joint to generate angular acceleration in the direction of the disturbance. This helps prevent falls and stabilize the upper body.

In this study, considering the pelvis tilt error from the vertical state as a destabilizing factor in human balance, the hip-ankle strategy is utilized during the stance phase to maintain balance. To implement this strategy, the distribution coefficients are assigned to the hip and ankle joints as,

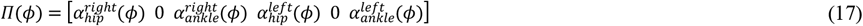

in which ϕ is the gait phase including “*Loading Response, Single Stance, Pre-Swing, Swing*”.

### Bio-inspired gait controller optimization

To optimize the unknown parameters of the gait controller, it is necessary to simulate the forward dynamics of the OpenSim model within a closed-loop platform. In present work, we developed a controllable interface between MATLAB and OpenSim software that enables us to interact with the OpenSim API (Application Programming Interface) using MATLAB commands. Specifically, within the interface, the gait controller script in MATLAB manipulates directly the Osim model by generating joint torques. Fig 1 displays the configuration of bio-inspired gait controller within OpenSim and MATLAB framework. Following equation represents the objective function of the optimization problem, which aims to minimize both the tracking error of the desired trajectory (***J***_*TE*_) and the mechanical energy (***J***_*ME*_).

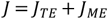

where,

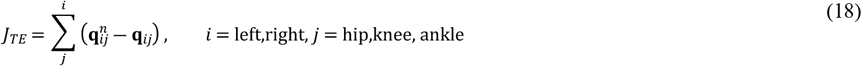

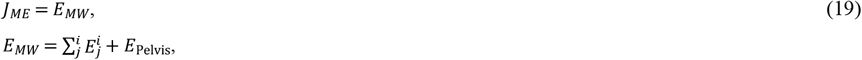

in which *E*_*j*_, and *E*_*Pelvis*_ is mechanical energy of lower limb joints and pelvis joint respectively. In the above equation, the mechanical energy is calculated by multiplying joint moment and joint angular velocity. The optimization constraint which is used to limit the range of optimal coefficients is defined as,

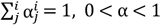

The optimization was performed on a computer with a 3.60 GHz Intel i5-8350U processor and 16 GB of RAM. The reported results were obtained using OpenSim release 4.3 and MATLAB release 2021b. The time simulation of a gait cycle (1.2 second) is approximately 2 minutes. Moreover, the optimization process of the gait balance control parameters took almost 24 hours to complete.

**Fig 1.**
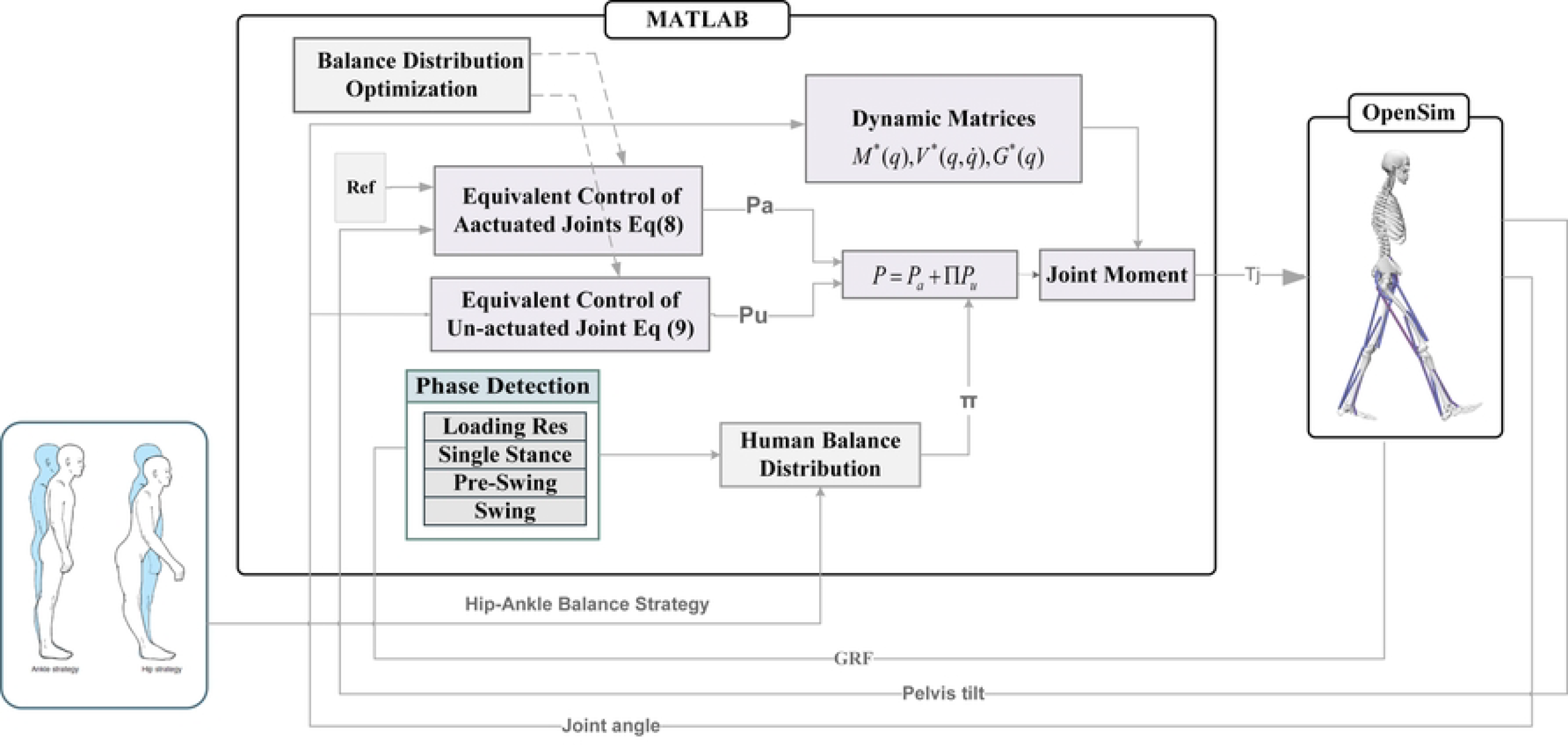
Schematic diagram of the Control Interface between MATLAB and OpenSim: The torque-driven skeletal human model is simulated using joint torques generated by a bio-inspired gait control script in MATLAB.

## Results & Discussion

Fig 2 displays the optimized balance distribution coefficients (BDC) and balance equivalent control torque (BEC) for the torque-driven model. The results are provided for a complete cycle of the right leg.

**Fig 2.**
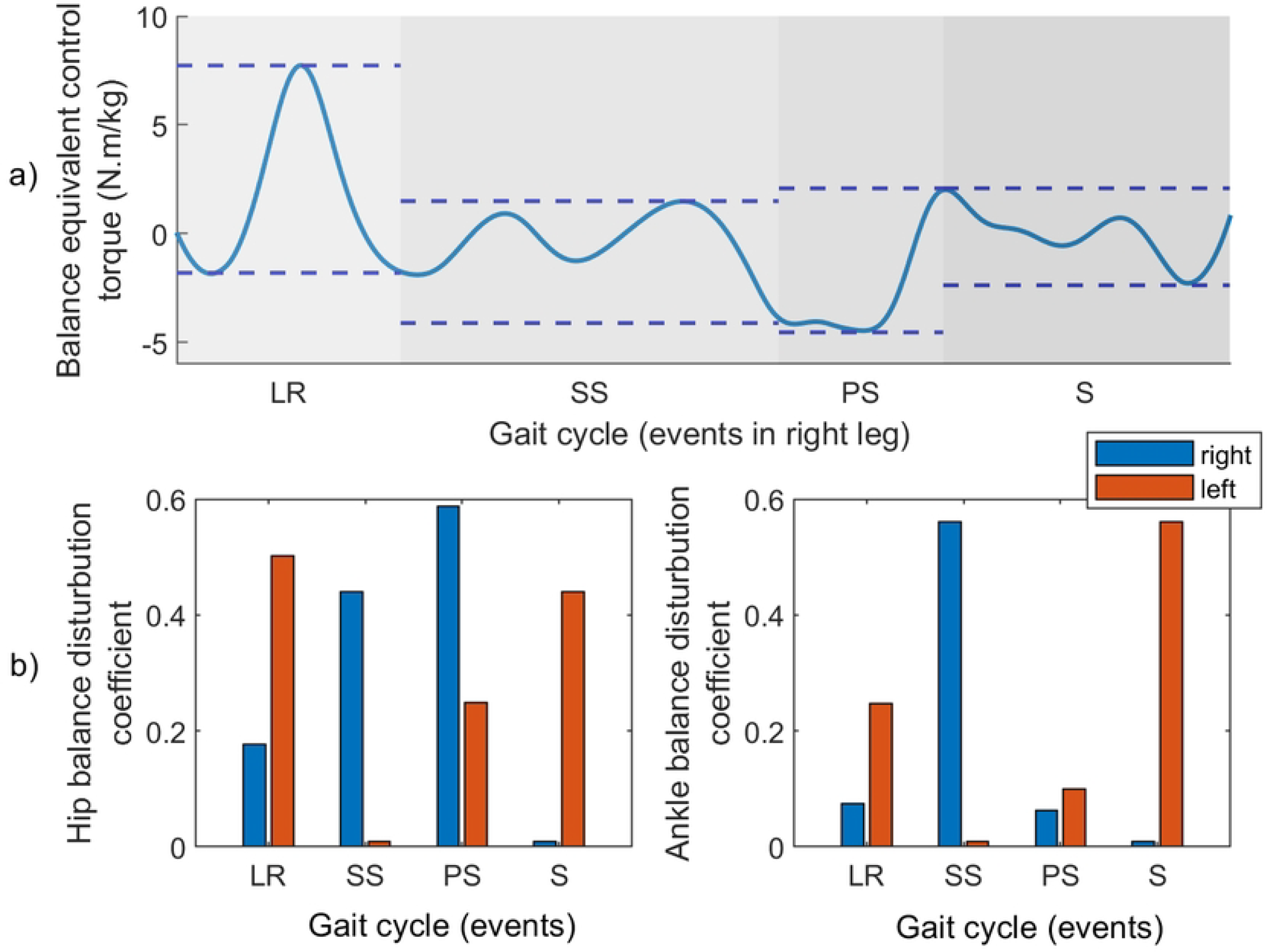
The optimized balance strategy versus the events in a complete gait cycle of the right leg; (a): The optimized Balance Equivalent Control Torque (BEC), (b): The optimized Balance Distribution Coefficients (BDC) of hip and ankle joints.

It is seen that the BEC undergoes changes during four gait events: loading response (LR), single stance (SS), pre-swing (PS), and swing (S). During the loading response, the BEC takes large values of up to 8 Nm/kg. In this phase, the BDC values for the right hip and ankle are 0.17 and 0.08, respectively, while for the left hip and ankle, they are 0.5 and 0.24, respectively. These BDC values indicate that, for large BEC values, more weight is assigned to the hip joint in both the right and left legs, which confirms the hip strategy. Obviously, when the right leg is in the loading response phase, the left hip, acting as the flexion limb, bears more body weight to maintain balance. As the result, in this phase, larger coefficients are assigned to the joints of the left leg. In the single stance phase, the average BEC takes some small values, indicating minimal disturbances on the upper body. As expected, there is an increase in the ankle BDC, reaching its maximum value of 0.58. This corresponds to the ankle strategy, where the ankle joint plays role in maintaining body balance when small disturbances act on the body. The maximum value of BEC changes from 2 N.m/kg in the single stance zone to 4 N.m/kg in the opposite direction in pre-swing phase, leading to significant body instability in this phase. Therefore, a significant increase in hip BDC is generated to overcome the instabilizing disturbances. Obviously, no coefficients were assigned to the right leg in the swing phase, indicating that the left ankle joint supports the body weight. Based on the alignment of simulation results with the biomechanical hip-ankle strategy, it can be concluded that the torque-driven control model exhibits a performance similar to the pattern of real human walking which attemts to maintain the human balance with the least energy consumption. In Fig 3, the gait cycles of the torque-driven model are simulated and compared to a normal human gait[38].

**Fig 3.**
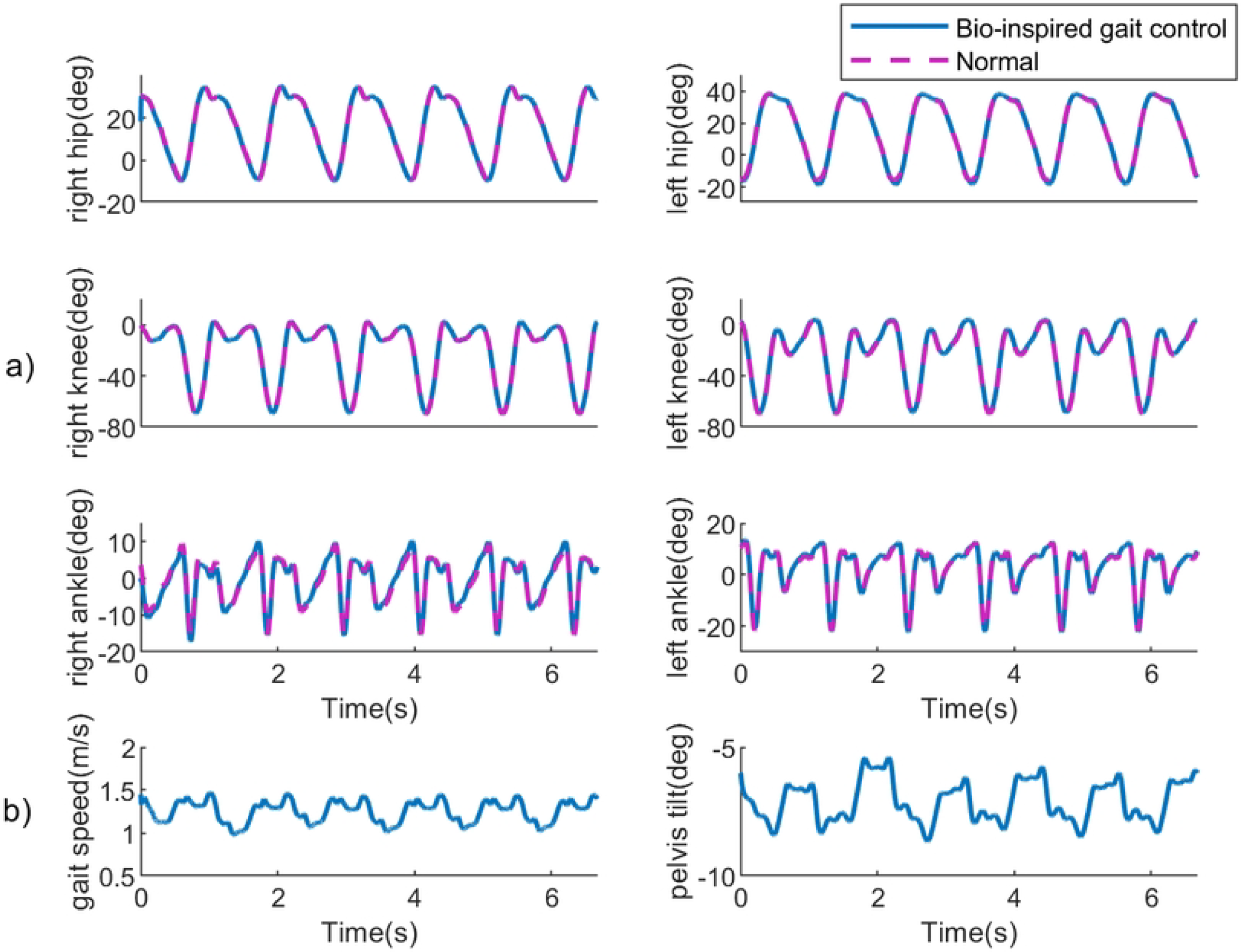
Kinematic results obtained from the optimized bio-inspired gait controller: a) Actuated joint trajectories vs. reference trajectories for hip, knee, and ankle in both right and left legs (reference trajectories are based on motion data of a normal human gait [39]), b) Un-actuated joint trajectories, including walking speed (m/s) and pelvis tilit (deg). Consistent walking speed and pelvis tilt demonsterate human balance during a five second walking simulation.

According to the figure, the joint angle patterns predicted by the bio-inspired gait controller closely resemble those observed in humans. Moreover, the limited changes in pelvic tilt and its vertical displacement indicate that the torque-driven model successfully prevents fall. Additionally, the consistent walking speed and angle of the unactuated pelvis joint demonstrates the maintenance of upper body balance throughout the walking process.

The Table 1 presents the joint angles RMSE of the model in comparison to the validated reference model. It’s evident from the table that the error in the hip and knee joint angles is less than 1 degree. However, the ankle angle exhibits an error greater than expected, possibly attributed to the contact force model.

**Table 1.** Root mean Squared error for the lower limb joint angles.

| Table 1. Root mean Squared error for the lower limb joint angles. |  |  |  |  |  |  |
| --- | --- | --- | --- | --- | --- | --- |
|  | Right Leg |  |  | Left Leg |  |  |
| RMSE (deg) | Hip | Knee | Ankle | Hip | Knee | Ankle |
|  | 0.5115 | 0.2557 | 1.0515 | 0.5642 | 0.2417 | 0.9588 |

Fig 4 provides a comparison between joint moments and ground reaction forces for torque-driven and musculoskeletal models. The model was used for comparison is the leg6dof9musc.osim model and simulated using the OpenSim in static optimization tool.

**Fig 4.**
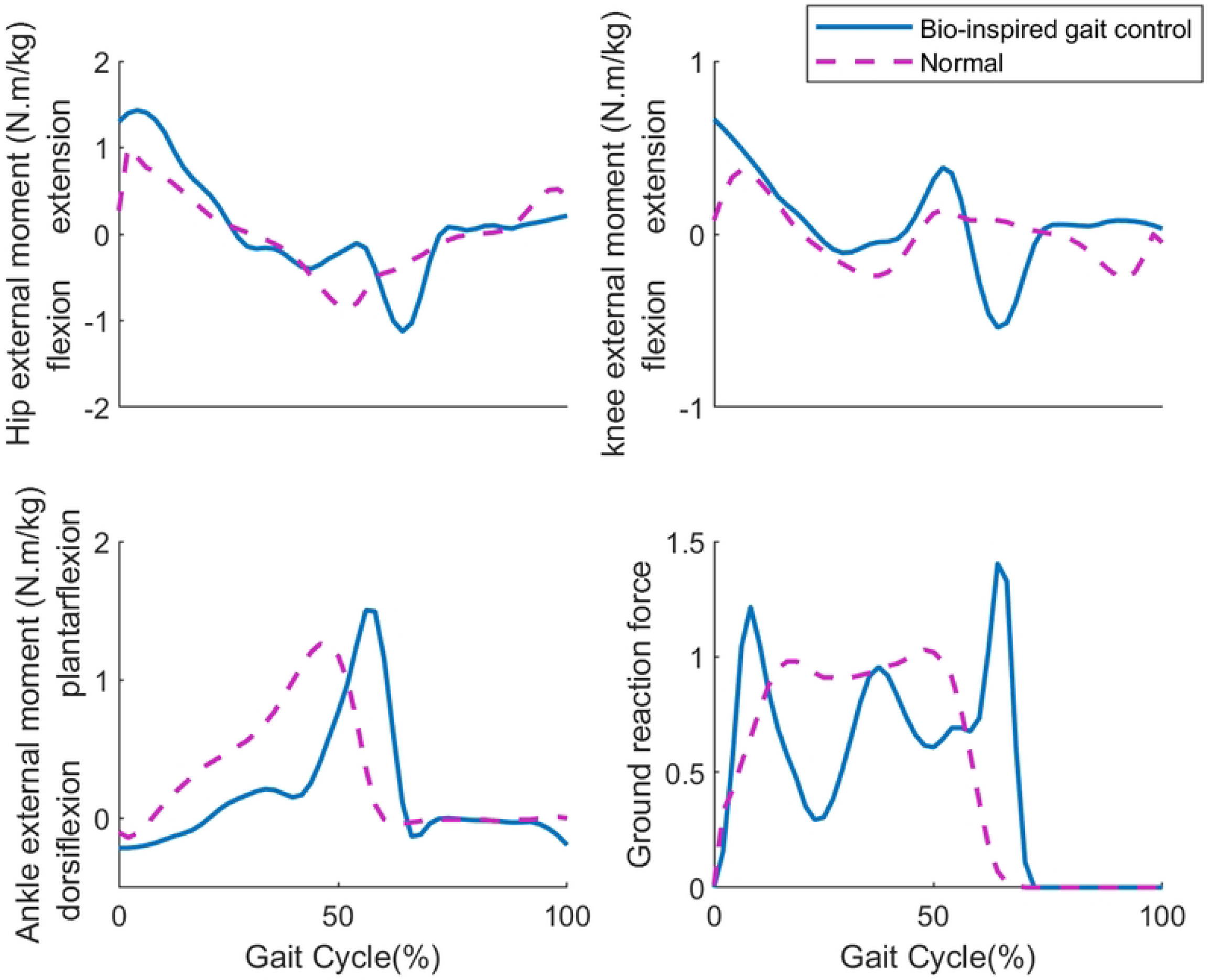
Comparison of joint moments (hip, knee and ankle) and ground reaction forces of right leg between the simulated torque-driven model and a muscle-driven model. The gait cycle starts at the heel-strike instant of the right leg. Data of the muscle-driven model is obtained by running the static optimiation for the leg6dof9musc.osim model in OpenSim.

Focusing on the hip joint of the muscle-driven model, a peak extension moment is observed during the loading response phase, while a peak flexion moment occurs in the pre-swing phase. The extension moment of the hip joint refers to the rotational force or torque generated when the hip joint moves into extension. This moment is primarily produced by the contraction of the hip extensor muscles, including the gluteus maximus, hamstrings (specifically the biceps femoris muscle), and adductor magnus. On the other hand, the flexion moment of the hip joint is primarily produced by the contraction of the hip flexor muscles, including the iliopsoas (composed of the iliacus and psoas major muscles), rectus femoris, and sartorius muscles. The comparison highlights the behavior of the torque-driven model, which approximately aligns with the musculoskeletal model.

When comparing knee joint patterns, it is observed that the torque-driven gait does not exhibit flexion during the pre-swing event. Additionally, a late knee flexion is observed in the swing phase, possibly due to the absence of weight assignment to the knee joint in this model. In terms of the ankle joint, a peak moment is identified in the plantarflexion moment. This moment allows the foot to push off the ground and propel the body forward. It is primarily generated by the contraction of the calf muscles, namely the gastrocnemius and soleus muscles, in the lower leg. The torque-driven model, influenced by the bio-inspired gait controller, effectively replicates this performance. Overall, Fig 4 demonstrates how both the torque-driven and musculoskeletal models exhibit similar joint moments, providing insights into the functional characteristics and muscle activations during various phases of gait. However, by improving the accuracy of phase detection, we can assign more distributed weights to the joints, resulting in more reliable simulation results.

The ground reaction force plays a crucial role in the study of human gait, providing valuable insights into the mechanics of walking. In the skeletal model, the foot-ground contact forces are modeled using the nonlinear Hunt-Crossley contact model on both the heel and toe of the foot, which is implemented in OpenSim. By comparing the ground reaction forces between the torque-driven model and the musculoskeletal model, we see a similar pattern of contact forces. However, there’s a significant reduction in force around the single stance phase. This reduction is particularly noticeable. The model only used two contact force models, specifically the toe and heel, and didn’t include a model for the middle of the foot. This absence of modeling resulted in inaccurate ground reaction force compared to the realistic data. Despite the drawback in force modeling, the results indicate that the torque-driven model closely approximates the interaction between the foot and the ground, similar to the observed behavior in actual human subjects.

In the next simulation, the robustness of the bio-inspired walking control is assessed by subjecting the model’s torso to disturbance forces of different amplitudes. These forces included low disturbance(30N), moderate disturbance(150N), and large disturbance (300N) applied horizontally, 20 centimeters above the center of mass of the torso.

Fig 5a illustrates the impact of these external disturbances on the deviations of the joint angles from their reference trajectories. The external forces were applied within the time duration from 1.7 to 1.8 seconds, as highlighted in gray. Additionally, Fig 5b presents changes in joint moment in response to external disturbance forces.

**Fig 5.**
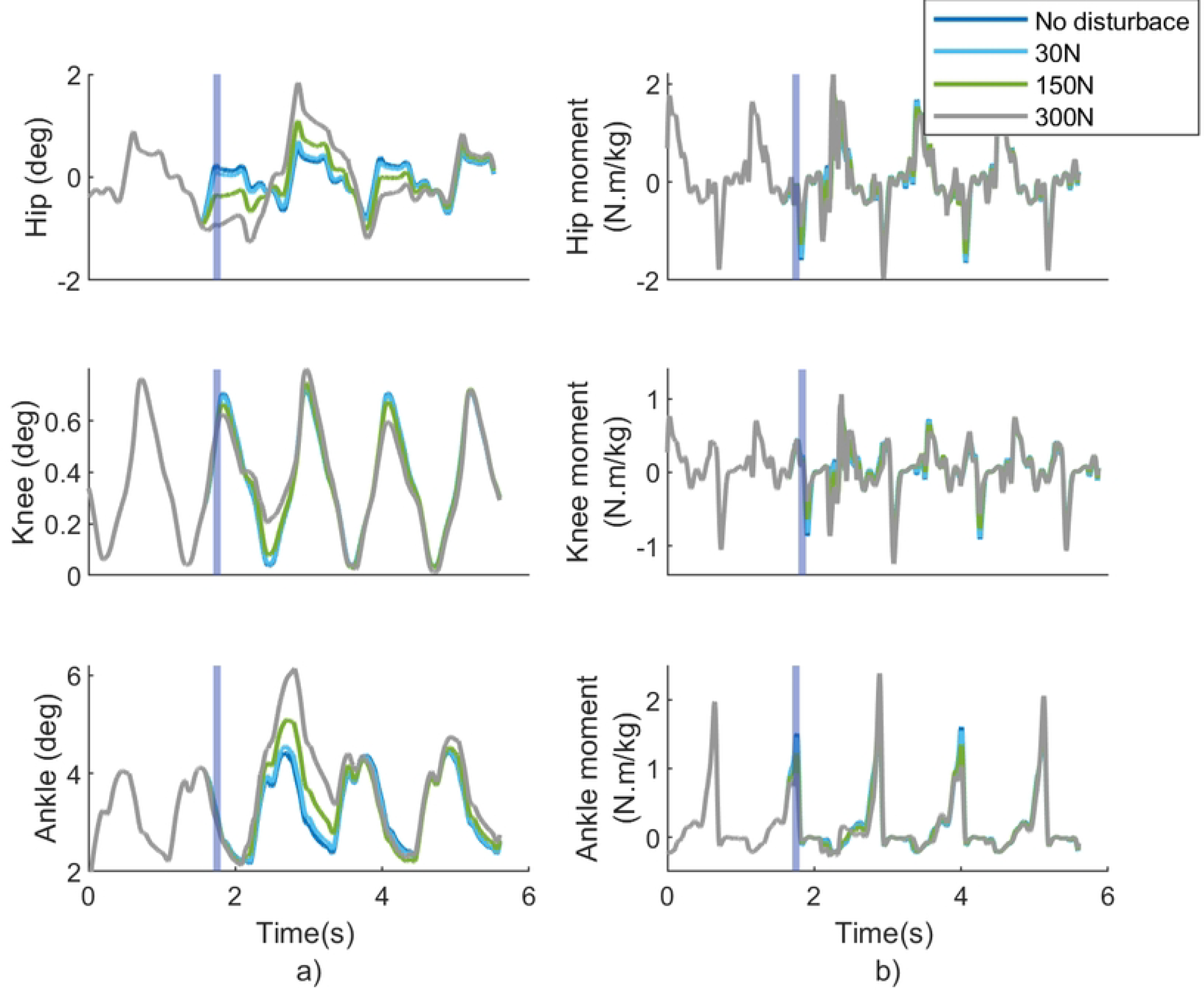
Performance of the torque-driven human model with bio-inspired gait controller against external disurbances. (a): Effect of the external disturbances on joint angle tracking, (b): Effect of the external disturbances on joint torques. Purpel areas indicates the time duration of disturbance force application.

In the case of a low-level disturbance, we observed slight alterations in the angle error and moments of the hip, knee, and ankle joints. However, as the disturbance level increases, more pronounced changes become evident in the hip and ankle joints. This observation demonstrates that these joints are generally more susceptible to external force disturbances compared to the knee. By applying large disturbance, the alterations in the hip and ankle joints becomes both more extensive and pronounced. Nevertheless, a significant deviation is seen in the hip angle error, which takes a relatively longer period of time to return to its original state compared to the ankle joint.

To enhance comprehension, we have graphed the R-Squared values for RMSE of joint angles in Fig 6a. Additionally, in Fig 6b, we have plotted the mean Pelvis tilt and mean walking speed versus different values of external forces. Fig 6a illustrates a clear linear correlation between joint angle RMSE and the increasing disturbance force. Notably, a relatively significant slope observed in the RMSE linear fitting for the hip and ankle joints which substantiates the significance of these two joints.

**Fig 6.**
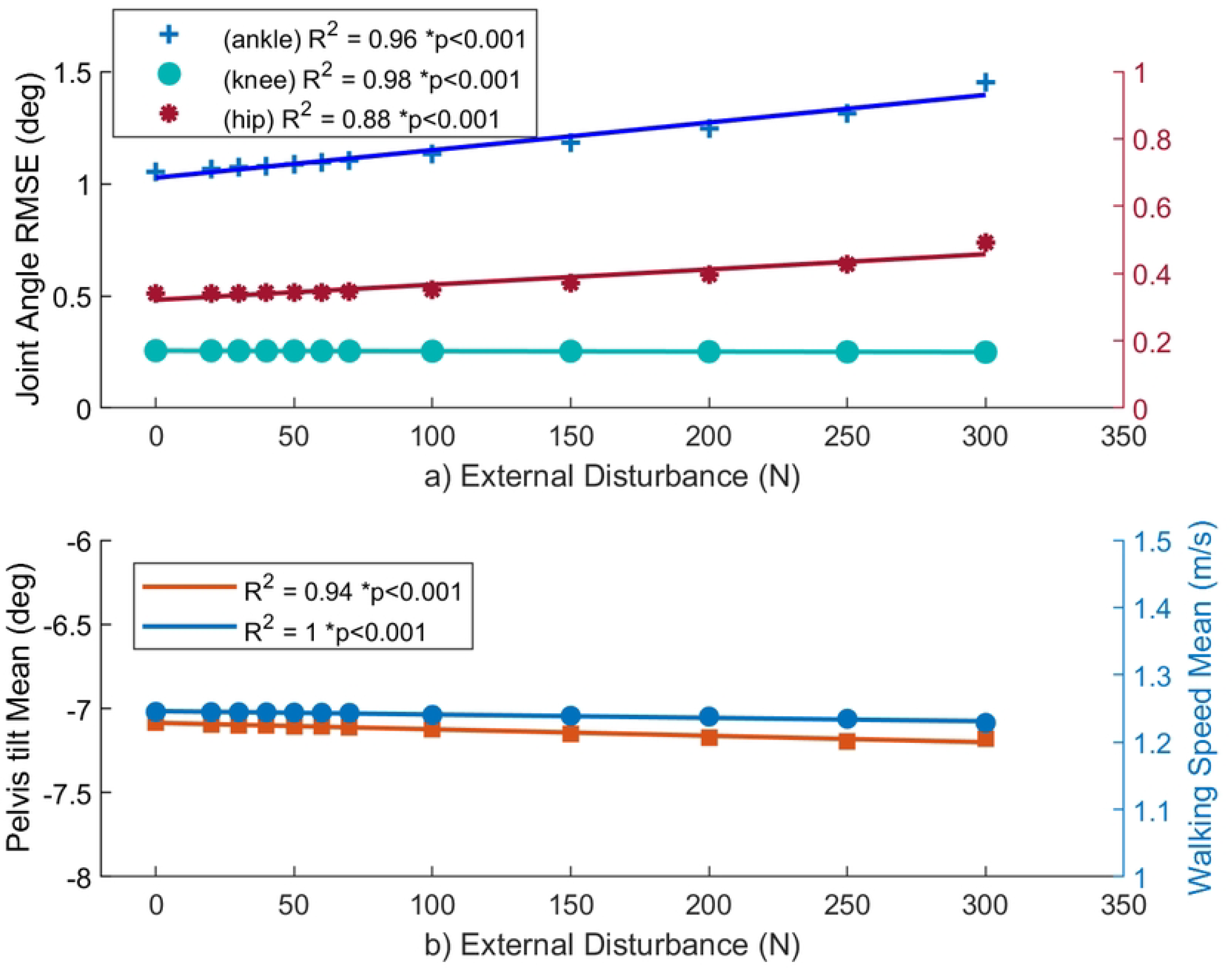
Correlation of motion kinematics with amplitude of the external disturbance, (a) correlation of root mean square error of torque-driven joint angles with external disturbance, (b) correlation of mean pelvis tilt angle and mean walking speed with external disturbance forces. P-value for all results is less than 0.001. R-Squared values are R^2^ =096, R^2^ =0.98,and R^2^ =0.88 for ankle, knee, and hip joint angles, and R^2^ =0.94 and R^2^ =1 for mean pelvis tilt and walking speed, respectively. This demonstrates the goodness of fit of the linear regressions. Increasing slopes of the linear regression for hip and ankle joints indicate relevant correlation of these joints with the external disturbance force. The almost horizontal lines in (b) indicate no correlation of tilt angle and walking speed with external disturbances, indicating the robustness of the balance controller.

This observation validates the efficacy of the hip-ankle strategy. Delving deeper, it’s worth noting that the slope of the ankle RMSE is slightly more than that of the hip. In fact, while both joints are more engaged during larger disturbances, the ankle joint assumes a more prominent role when dealing with lower levels of disturbance. Analyzing Fig 6b, the low slope exhibited by the mean value of unactuated data (including both mean pelvis tilt (on the left Y-axis) and mean walking speed (on the right Y-axis), indicates a weak correlation between these variables and external forces. This association highlights the successful performance of the bio-inspired walking controller in maintaining the human balance without affecting the overall characteristics of the motion.

## Conclusion

In this paper, we introduced a bio-inspired gait controller for a torque-driven skeletal model to achieve a stable walking with joint trajectory tracking. Since the model is under-actuated and requires a balance maintenance approach, we wxtended the hip-ankle strategy from the usual cases of upright standing to the case of walking. First, a “*Balance Equivalent Control Torque*” is calculated using a model-based approach to keep the upper body in upright position. Then, this torque is distributed to the hip and ankle joints of both legs, in proportion to some “*Balance Distribution Coefficients*”. This strategy was applied during the loading response, single stance, and pre-swing phases. To adjust the controller parameters, we established a closed-loop interface between OpenSim and MATLAB for simulating the forward dynamic model. The optimization results confirmed that the torque-driven skeletal model with the proposed controller, exhibits human-like behavior in intrinsically maintaining upper body stability during a walking gait. For instance, when the average balance equivalent control (BEC) is low in the single stance phase, the ankle joint coefficient increases to compensate for the instability. Similarly, if the average BEC is high, the hip joint acts as a balancer. Furthermore, in terms of joint moments, we observed a similar performance between the hip and ankle joints during the loading response and pre-swing phases. This similarity further supports the effectiveness of the implemented hip-ankle strategy and its ability to replicate human-like walking characteristics. In addition, to assess the robustness of the controller, we exerted a range of horizontal forces on the torso as external disturbances. The simulation results proved the balance recovery performance of the proposed biomechanical gait controller. Overall, our findings demonstrate that the combination of the bio-inspired gait controller and the joint-space model yields results that closely resemble human gait, particularly in terms of maintaining stability and replicating joint moments during specific gait events.

## Author Contribution

**Conceptualization**: Samane Amini, Iman Kardan, Alireza Akbarzadeh

**Data curation:** Samane Amini

**Investigation:** Samane Amini, Iman Kardan, Ajay Seth, Alireza Akbarzadeh

**Methodology:** Samane Amini, Iman Kardan, Ajay Seth, Alireza Akbarzadeh

**Writing – original draft:** Samane Amini

**Writing – review & editing:** Iman Kardan, Ajay Seth

## Notes

### Competing Interest Statement

The authors have declared no competing interest.

## Reference

[1] S. A. Ghafari, A. Meghdari, and G. Vossoughi, “Forward dynamics simulation of human walking employing an iterative feedback tuning approach,” Proceedings of the Institution of Mechanical Engineers, Part I: Journal of Systems and Control Engineering, vol. 223, no. 3, pp. 289–297, 2009.

[2] B. Abdikadirova, M. Price, W. Hoogkamer, and M. E. Huber, “Forward simulations of walking on a variable surface-impedance treadmill: A comparison of two methods,” bioRxiv, p. 2021.10. 11.463993, 2021.

[3] N. Waterval et al., “Validation of forward simulations to predict the effects of bilateral plantarflexor weakness on gait,” Gait & Posture, vol. 87, pp. 33–42, 2021.

[4] T. Nomura, K. Kawa, Y. Suzuki, M. Nakanishi, and T. Yamasaki, “Dynamic stability and phase resetting during biped gait,” Chaos: An Interdisciplinary Journal of Nonlinear Science, vol. 19, no. 2, 2009.

[5] L. Liu, J. L. Cooper, and D. H. Ballard, “Computational Modeling: Human Dynamic Model,” (in English), Frontiers in Neurorobotics, Methods vol. 15-2021,September-24 2021, doi: 10.3389/fnbot.2021.723428.

[6] R. M. Alexander, “Simple Models of Human Movement,” Applied Mechanics Reviews, vol. 48, no. 8, pp. 461–470, 1995, doi: 10.1115/1.3005107.

[7] J. Laczkó, “Modeling of human movements, neuroprostheses,” Ideggyogyaszati Szemle, vol. 64, no. 7-8, pp. 229–233, 2011.

[8] H. Hemami and V. C. Jaswa, “On a three-link model of the dynamics of standing up and sitting down,” IEEE Transactions on Systems, Man, and Cybernetics, vol. 8, no. 2, pp. 115–120, 197.8

[9] J. L. Hicks, T. K. Uchida, A. Seth, A. Rajagopal, and S. L. Delp, “Is my model good enough? Best practices for verification and validation of musculoskeletal models and simulations of movement,” Journal of biomechanical engineering, vol. 137, no. 2,p. 020905, 2015.

[10] S. L. Delp et al., “OpenSim: open-source software to create and analyze dynamic simulations of movement,” IEEE transactions on biomedical engineering, vol. 54, no. 11, pp. 1940–1950, 2007.

[11] A. Seth et al., “OpenSim: Simulating musculoskeletal dynamics and neuromuscular control to study human and animal movement,” PLoS computational biology, vol. 14, no. 7, p. e1006223, 2018.

[12] M. Benallegue and J.-P. Laumond, “Bipedal locomotion: a continuous tradeoff between robustness and energy-efficiency,” Biomechanics of Anthropomorphic Systems, pp. 263–279, 2019.

[13] S. Gupta and A. Kumar, “A brief review of dynamics and control of underactuated biped robots,” Advanced Robotics, vol. 31, no. 12, pp. 607–623, 2017.

[14] S. Park and J .Oh, “Real-time continuous ZMP pattern generation of a humanoid robot using an analytic method based on capture point,” Advanced Robotics, vol. 33, no. 1, pp. 33–48, 2019.

[15] J. Liu and O. Urbann, “Bipedal walking with dynamic balance that involves three-dimensional upper body motion,” Robotics and Autonomous Systems, vol. 77, pp. 39–54, 2016.

[16] H. F. Al-Shuka, B. Corves, W.-H. Zhu, and B. Vanderborght, “Multi-level control of zero-moment point-based humanoid biped robots: a review,” Robotica, vol. 34,no. 11, pp. 2440–2466, 2016.

[17] Z. Wang, B. He, Y. Zhou, T. Yuan, S. Xu, and M. Shao, “An experimental analysis of stability in human walking,” Journal of Bionic Engineering, vol. 15, pp. 827–838, 2018.

[18] T. Yamasaki, T. Nomura, and S. Sato, “Possible functional roles of phase resetting during walking,” Biological cybernetics, vol. 88, pp. 468–496, 2003.

[19] A. Macchietto, V. Zordan, and C. R. Shelton, “Momentum control for balance,” in ACM SIGGRAPH 2009 papers, 2009, pp. 1–8.

[20] T. Yamasaki, T. Nomura, and S. Sato, “Phase reset and dynamic stability during human gait,” Biosystems, vol. 71, no. 1-2, pp. 221–232, 2003.

[21] S. Aoi, N. Ogihara, Y. Sugimoto, and K. Tsuchiya, “Simulating adaptive human bipedal locomotion based on phase resetting using foot-contact information,” Advanced Robotics, vol. 22, no. 15, pp. 1697–1713, 2008.

[22] B. Ugurlu et al., “Variable ankle stiffness improves balance control: Experiments on a bipedal exoskeleton,” IEEE/ASME Transactions on mechatronics, vol. 21, no. 1,pp. 79–87, 2015.

[23] N. Hogan, “The mechanics of multi-joint posture and movement control,” Biological cybernetics, vol. 52, no. 5, pp. 315–331, 1985.

[24] H.-O. Lim, S. A. Setiawan, and A. Takanishi, “Position-based impedance control of a biped humanoid robot,” Advanced Robotics, vol. 18, no. 4, pp. 415–435, 2004.

[25] A. Bottaro, Y. Yasutake, T. Nomura, M. Casadio, and P. Morasso, “Bounded stability of the quiet standing posture: an intermittent control model,” Human movement science, vol. 27, no. 3,pp. 473–495, 2008.

[26] I. D. Loram, H. Gollee, M. Lakie, and P. J. Gawthrop, “Human control of an inverted pendulum: is continuous control necessary? Is intermittent control effective? Is intermittent control physiological?,” The Journal of physiology,vol. 589, no. 2, pp. 307–324, 2011.

[27] Y. Suzuki, T. Nomura, M. Casadio, and P. Morasso, “Intermittent control with ankle, hip, and mixed strategies during quiet standing: a theoretical proposal based on a double inverted pendulum model,” Journal of theoretical biology, vol. 310, pp. 55–79, 2012.

[28] J. L. Cabrera and J. G. Milton, “On-off intermittency in a human balancing task,” Physical Review Letters, vol. 89, no. 15, p. 158702, 2002.

[29] V. T. Inman, “Human locomotion,” Canadian Medical Association Journal, vol. 94, no. 20, p. 1047, 1966.

[30] M. Zarrugh, F. Todd, and H. Ralston, “Optimization of energy expenditure during level walking,” European journal of applied physiology and occupational physiology, vol. 33, pp. 293–306, 1974.

[31] N. Seethapathi and M. Srinivasan, “The metabolic cost of changing walking speeds is significant, implies lower optimal speeds for shorter distances, and increases daily energy estimates,” Biology letters, vol. 11, no. 9, p. 20150486, 2015.

[32] J. A. Schrack, E. M. Simonsick, and L. Ferrucci, “The relationship of the energetic cost of slow walking and peak energy expenditure to gait speed in mid-to-late life,” American journal of physical medicine & rehabilitation/Association of Academic Physiatrists, vol. 92, no,1 .p. 28, 2013.

[33] P. Morasso, “Integrating ankle and hip strategies for the stabilization of upright standing: An intermittent control model,” Frontiers in Computational Neuroscience, vol. 16, 2022.

[34] G. M. Blenkinsop, M. T. Pain, and M. J. Hiley “,Balance control strategies during perturbed and unperturbed balance in standing and handstand,” Royal Society open science, vol. 4, no. 7, p. 161018, 2017.

[35] C. L. Dembia, N. A. Bianco, A. Falisse, J. L. Hicks, and S. L. Delp, “Opensim moco: Musculoskeletal optimal control,” PLOS Computational Biology, vol. 16, no. 12, p. e1008493, 2020.

[36] S.-G. Lee, “Nonlinear feedback control of underactuated mechanical systems,” in Nonlinear Systems-Design, Analysis, Estimation and Control: IntechOpen, 2016.

[37] D. N. Nenchev and A. Nishio, “Ankle and hip strategies for balance recovery of a biped subjected to an impact,” Robotica, vol. 26, no. 5, pp. 643–653, 2008.

[38] G. Bovi, M. Rabuffetti, P. Mazzoleni, and M. Ferrarin, “A multiple-task gait analysis approach: kinematic, kinetic and EMG reference data for healthy young and adult subjects,” Gait & posture, vol. 33, no. 1, pp. 6–13, 2011.

